# Device for Axon - Cancer cell Interaction Testing in 2D & 3D: DACIT

**DOI:** 10.1101/2024.02.26.582218

**Authors:** Ines Velazquez-Quesada, Vahid Alizadeh, Kara Allison, Elizaveta Belova, Svetllana Kallogjerovic, Natasha Hesketh, Xiaotian Zhang, Gareth Thomas, Erkan Tuzel, Bojana Gligorijevic

**Author notes:** Correspondence: Bojana Gligorijevic.

## Abstract

There is increasing interest in studying the role of peripheral innervation in tumor growth and metastasis. However, *in vitro* studies of interactions between cancer cells and axonal projections are technically challenging. To address this, we have developed a microfluidic Device for Axon-Cancer cell Interaction Testing in 2D and 3D (DACIT). We show that DACIT successfully separates neuronal soma from the axons and cancer cells into two compartments, which can be exposed to similar, or different growth conditions, depending on the experimental needs. We compare neoaxonogenesis using either the PC-12 cell line or primary embryonic or adult sensory neurons, demonstrating superior neurite growth in primary cells. Additionally, we show that DACIT can accommodate assessing growth and 3D invasion of tumor spheroids, due to its unique height profile. Hence, DACIT can be used to analyze cancer cell interactions with axons in most typical cell biology assays such as proliferation, invasion, and calcium activity which we demonstrate on examples of imaging transients in GCaMP6-labeled neurons, invadopodia assay, and 3D cancer spheroid invasion.

## Introduction

While the nerve infiltration into tumors was identified more than 100 years ago, it wasn’t until recently that neoaxonogenesis has been recognized as one of the hallmarks of cancer (Baraldi et al., 2022; Hanahan & Monje, 2023). Even now, cancer cell-neuron interactions are mostly studied in the central nervous system, while the interactions between neurons and cancer cells in the peripheral organs remain generally understudied. Few recent studies have started demonstrating the impact of the autonomic or somatic peripheral nervous system in tumor growth and metastasis, including pancreatic, melanoma and breast cancer (Kappos et al., 2018; Le et al., 2021; Peterson et al., 2015; Prazeres et al., 2020). These observations generated an increased interest in understanding the molecular and cellular mechanisms that drive interactions between peripheral neurons and cancer cells and developing appropriate technical solutions for such studies.

Several technical reasons can explain this knowledge gap. First, mammals have a unique anatomy where the peripheral neurons’ somas (cell bodies) are located in ganglia distributed along the body: close to the cranial nerve, in the vicinity of the vertebral column, or the abdominal cavity. Consequently, innervation of peripheral organs is achieved via axons and axon bundles (fascicles). In humans, distance between the ganglion and the organ, or tumor, that the ganglion innervates can be longer than one meter. Mimicking such cellular compartmentalization in tissue culture-based experiments is important to maintain anatomical similarity and avoid experimental artifacts that can occur in simple co-culture settings. Neurotoxicity of the glutamate, which appears at high concentrations in the fetal bovine serum used for culturing most cancer cell lines is one classical example of such experimental differences (Brewer, 1998). Second, currently available models of peripheral neurons do not reflect the complexity and diversity of the normal peripheral nervous system (Chrysostomidou et al., 2021; Jones et al., 2018; Li et al., 2022). Consequently, most studies use embryonic or adult primary culture from rats or mice to study neuron behavior. While a few recent studies have reported successful differentiation of induced pluripotent stem cells into sensory neurons (Deng et al., 2023; Hiranuma et al., 2024b; Labau et al., 2022), this approach is not yet widely utilized. However, the use of primary culture limits the number of neurons that can be obtained at a given time, and also introduces the issue of the presence of non-neuronal cells, such as Schwann cells, which affect cancer cell behavior (Deborde et al., 2016; Deborde & Wong, 2022) but can be difficult to eliminate without affecting the health of neurons.

To provide the compartmentalization between soma and axons, Campenot developed a chamber consisting of a collagen-coated Petri dish with scratches to guide axon growth (Campenot, 1977). Recently, principles of microfluidics were added to the Campenot chamber, with several variations. Since then, the progress of microfabrication techniques and the inspiration of the original concept of the Campenot chamber triggered the development of a series of compartmentalized microfluidic devices to analyze anything from axonal regeneration (Park et al., 2006; Taylor et al., 2005a) to cellular and biochemical assays or drug screening (Neto et al., 2016) in 2D cultures.

In this paper, we focused on developing a microfluidic chamber which can allow analysis of both 2D and 3D cell cultures. The 3D culture represents a physiologically relevant alternative to animal models, as it better recapitulates *in vivo* cell behaviors, including proliferation, morphology, adhesion, motility, etc.(Evans & Teicher, 2017; Shamir & Ewald, 2014). Here, we present a novel Device for Axon Cancer cell Interaction Testing in 2D and 3D (DACIT). DACIT allows monitoring of the interactions between axons and cancer cells growing on thin layers of extracellular matrix (ECM) in 2D, or spheroids embedded in the 3D ECM. DACIT can be used to expose neuronal soma and axons to different, or similar, media conditions, using compartmentalization or diffusion/mixing of media, respectively. Moreover, DACIT allows genetic and chemical treatments and high-resolution, time-lapse imaging.

## Methods

### Design and fabrication of DACIT mask and master

DACIT was designed utilizing LayoutEditor software and consists of two layers. The first layer contains microgrooves which connect left and right macrochannels; the second layer was used for generating the macrochannels, which house neurons on the left, and axons and cancer cells on the right side. Microgrooves and macrochannel chrome masks (5" x 5" x 0.090" Quartz Plate, QZ) were produced by Compugraphics (Waterbury, CT). The details of both these steps are as follows:

### Microgroove layer fabrication using SU-8 photoresist

To fabricate a very thin layer with strong attachment properties, we used SU8-3000 series photoresist (KAM-SU-8-3000, Kayaku Advanced Materials Inc., Waterborough, MA). SU8-3050 stock was diluted to the viscosity of SU8-3005 using the formula m1v1=m1v2 and spin-coated. The wafer was soft-baked for 2 min at 95°C (1st bake, **Figure 2A**, top row) and exposed to 100 mJ/cm^2^ of UV light in the mask aligner (ABM 3000HR Mask Aligner, ABM-USA, San Jose, CA). The exposed layer went through two post-exposure bakes: one for 1 min at 65°C and one for 1 min at 95°C. The layer was developed after cooling down, using SU-8 developer (Kayaku Advanced Materials Inc., Waterborough, MA).

### Alignment and addition of macrochannel layer to the microgrooves

A chrome mask (5" x 5" x 0.090" QZ) or a transparency 10 000 dpi film (ArtNet Pro, San Jose, CA) were used. To fabricate a thick layer, SU8-2100 stock was spin-coated on the wafer without dilution, with the alignment marks covered using duct tape. Duct tape was removed before the 2^nd^ soft bake, which was done at 95°C for 10 min. Next, a second layer of SU8-2100 was added and spin-coated on top of the first, to increase the channel height. The 3^rd^ soft bake at 95°C for 30 min was done, and finally the wafer with a thick layer of baked SU8-2100 was placed in mask aligner (ABM 3000HR Mask Aligner, ABM-USA, San Jose, CA). The two alignment marks on photomask and wafer were carefully aligned utilizing x, y, and 𝜃 (angular orientation) knobs, and the layer was exposed to 300 mJ/cm^2^ of UV light. The exposed layer went through post-baking for 5 min at 65°C and 20 min at 95°C. After cooling down, the wafer with SU8 was developed for 30 min with constant swirling in SU8 developer. Finally, the master was washed with acetone, isopropanol, and water in order and dried with air flow. The cleaned master was hard-baked at 150°C for 10 min for thermal stability (**Figure 2A**, middle and bottom rows).

### DACIT device fabrication

DACITs are made from polydimethylsiloxane (PDMS) using the silicone elastomer (Sylgard 184, Sigma-Aldrich, Burlington, MA). Briefly, the elastomer base and the curing agent (Electron Microscopy Sciences, Cat#24236-10) were mixed at a 1:10 ratio, poured on the SU-8 master, and cured overnight at 65°C. The next day, the cured PDMS was peeled off, individual DACITs were cut out using scalpel and wells were punched out using 6 mm diameter tissue punches (Robbins instruments, Cat#RBP-20). Individual DACIT devices were plasma bonded to clean 24 x 24 mm coverslips (Globe Scientific Inc, Cat #1405-10). DACITs were sterilized, first by a wash with 70% ethanol, and then by a 15 min exposure to the biosafety cabinet UV light. Next, DACITs were coated with 50 μg/ml Poly-L-lysine overnight, followed by 30 min incubation with 1:10 Matrigel® Basement Membrane Matrix (Corning, Cat#356234) diluted in PBS.

### Institutional permissions

All experimental procedures were approved by the Institutional Animal Care and Use Committee of Temple University.

### Sensory neuron dissection and dissociation

In adult mice, DRG dissection and dissociation were performed as described previously (Theresa Heinrich*, 2016). Briefly, mice were sacrificed, the spinal cord was removed, and maximum number of DRGs (∼30) per mouse were recovered. Recovered DRGs were digested with 5 mg/ml Collagenase P (Sigma-Aldrich, Cat#11249002001) for 45 min at 37°C, followed by a 5 min digestion by trypsin/EDTA at 37°C. Trypsin was inactivated with 10% horse serum (Atlanta, Cat#S12150 in a Neurobasal medium (Gibco, Cat# 21103049), and cells were disaggregated using a polished pipet. Cells suspension was filtered through a 40 µm cell mesh strainer (Corning, Cat#431750) to remove cell clusters and centrifugated through a 10% bovine serum albumin (BSA, Sigma-Aldrich, Cat# Cat#A4503) layer, to remove debris and non-neuronal cells. Cells were concentrated by centrifugation, resuspended at 125,000 cells/ml in adult DRG media and plated in the DACITs.

In rat embryos, DRG dissection and dissociation was done at E16. Embryos were removed and DRGs were collected and placed on cold HBSS. Embryonic rat DRGs were first digested with Collagenase (10kU/ml) for 15 min at 37°C. Trypsin-EDTA was then added and DRGs were digested for a further 15 min at 37°C. Trypsin was inactivated with FBS and washed out by centrifugation. Dissociated DRGs were triturated in Neurobasal media using pipet tips (P1000 and then P200) before cell counting. Cells were resuspended in embryonic DRG media and loaded into DACIT as described.

### Cell culture

All cells were maintained at 37°C with 5% CO_2_ in a medium with 100 U/ml of Penicillin-Streptomycin (Gibco, Cat#15140122). Murine mammary cancer cell line 4T1 was grown in cancer media, consisting of DMEM (Gibco, Cat#10-013-CV) with the addition of 10% fetal bovine serum (FBS, Atlanta Biologicals, Cat#S11550). Neuronal PC12 cell line was maintained in DMEM medium with 7.5% FBS, and 7.5% horse serum and differentiated with adding 25 ng/ml of Neural Growth factor (NGF, Invitrogen, Cat#13257019). The primary culture of adult neurons was maintained in adult DRG media, consisting of neurobasal medium (Gibco, Cat# 21103049) complemented with B27 (Gibco, Cat# 17504044), and Penicillin-Streptomycin. Primary culture of embryonic sensory neurons was maintained in embryonic DRG media, consisting of neurobasal media supplemented with B27, Glutamax (Gibco, Cat#35050061), FdU (Sigma-Aldrich, Cat#PHR2589) and 25 ng/ml of NGF.

Cancer cell spheroids were prepared using the hanging drop method as described previously (Perrin et al., 2021, Perrin et al 2022). Briefly, 1250 4T1 cells were placed in 1 ml of complete DMEM media (Gibco, Cat# 11965118) containing 4.8 mg/ml methylcellulose (Sigma-Aldrich, Cat# M6385) and 20 µg/ml Nutragen (Advanced Biomatrix, Cat#5010-D). Drops containing 50 cancer cells were placed on the lid of a cell culture dish and incubated 24-36 h at 37°C with 5% CO_2_. Properly formed spheroids were selected by morphology and loaded in DACIT.

### DACIT coating with ECM and loading with cells

#### ECM coating for the 2D invadopodia-mediated degradation assays

Prior to loading with neurons and cancer cells, the axonal compartment was coated with 0.2% fluorescently labeled gelatin, as described in a previous study (Perrin et al., 2022). Briefly, the DACIT’s axonal compartment was treated sequentially with 1N HCl (Fisher Scientific, Cat#SA48-1), 50 µg/ml poly-L-lysine (Sigma, Cat#P8920), and 0.2% labeled gelatin. Coated devices were cross-linked with 0.2% glutaraldehyde (Sigma, Cat#G5882) and quenched with 50mM Glycine (ThermoScientific, Cat#J62407.22). To promote axonal growth, gelatin layer was further coated with 50 µg/ml poly-L-lysine (Sigma, Cat#P8920) for 20 min at room temperature, washed twice with PBS, and incubated with 0.15 mg/ml laminin (Sigma, Cat#L2020) for 20 min at room temperature.

### Loading with neurons

10,000 DRG cells resuspended in 10 µl of media were loaded into the neuronal compartment of DACIT. After allowing 20 min for cell attachment, 190 µl or 150 µl of media were added in the neuronal or axonal compartment, respectively. The following day, fresh media containing 50 ng/ml of NGF was added into the axonal compartment, equalizing the volumes of liquid present in the two compartments such that NGF diffuses across the microgrooves. At 3 days *in vitro* (DIV), media was removed from the axonal compartment, and the compartment was washed twice with HBSS without Ca^2+^, Mg^2+^ (Gibco, Cat#14170161).

### Loading with cancer cells

For 2D assays, 7,500 cancer cells in 10 µl of media were loaded into the axonal compartment of DACIT. After 30 min, 50 µl of adult DRG media was added to the neuronal compartment and 140 µl of cancer cell media was added to the axonal compartment. Cells were incubated at 37°C with 5% CO_2_ until fixation, replenishing media every 48h to avoid DACIT drying out.

For 3D assays, 50-cell spheroids were prepared using 4T1 cells. Spheroids were checked for round morphology using light microscope, washed 3 times with warm DMEM, and embedded in 2 mg/ml of collagen I (Corning, Cat#354249) with 20% of Matrigel (Corning, Cat#356234) on ice to avoid polymerization. Spheroids were loaded into the axonal compartment, and DACITs were incubated at 37°C with 5% CO2 until analysis.

### Transduction of DRG Cells in DACIT

The soma and axonal compartments of DACIT were coated with 50 µg/mL Poly-L-lysine, followed by a 1:20 dilution of Matrigel in PBS, as previously described. Next, 10,000 DRG cells were plated into the axonal compartment in complete Neurobasal medium. At 2 DIV, AAV5- syn-GCaMP6f (Addgene, Cat#100837-AAV5) was added to the neuronal compartment at a multiplicity of infection (MOI) of 10^5^ viral genomes per cell. Expression of GCaMP6f was monitored daily using a Nikon Eclipse Ti2 microscope. At DIV5, GCaMP6f-expressing cells were observed, and 1,000 4T1 breast cancer cells were plated into the axonal compartment of DACIT.

### Immunofluorescence

Immunofluorescence of neurons and spheroids in DACIT, was previously described (Perrin et al., 2021). Briefly, media was removed, and the cells were washed with PBS. Cells were incubated with a fixing/permeabilizing solution (4% PFA with 0.5% Triton X-100 in PBS) for 10 min at room temperature. Solution was removed, and cells were incubated with fixing solution (4% PFA in PBS) for 20 min at room temperature. After fixation, cells were washed thrice with washing solution (PBS containing 0.05% Tween-20), and unspecific sites were blocked with blocking solution (1% FBS, 1% BSA in PBS) for 1 hour at room temperature. Blocking solution was removed, and cells were incubated overnight at 4°C with the primary antibodies diluted in the blocking solution. The next day, the primary antibody was removed, and the cells were washed thrice with washing solution for 10 min, at room temperature. Then, cells were incubated for 1 h at room temperature with the secondary antibodies diluted in the blocking solution. After incubation, cells were washed thrice with the washing solution for 10 min, at room temperature. DACITs were stored at 4°C, in PBS-containing chambers, to avoid evaporation and preserve high humidity in the device.

### Imaging and axon quantification

To mount and image DACITs, custom-designed holders for individual DACITs or 6 DACITs were 3D printed using ABS, or CNC-machined using aluminum. Imaging was done using the confocal microscope (Olympus FV12000MPE), with UPLSAPO10X2 and UPLSAPO30XS objectives. For live imaging, DACITs were placed on sterilized aluminum holders and kept at 37°C with 5% CO_2_ and 95% humidity using an environmental chamber (STXG-WELSX-SET, Tokai Hit, Japan). To avoid axon damage, imaging was performed using low-laser power. GCaMP6f imaging was conducted using confocal microscope (FV3000, Olympus) equipped with a resonant scanner and a 30× oil-immersion objective (UPLSAPO30XS, 1.05 NA, Olympus), at a frame rate of 14 Hz.

The length of the axons was tracked using maximum intensity projection of z-stacks in NeuronJ plugin (Meijering et al., 2004) in Fiji, ImageJ2 2.9.0/1.53t. Axons were measured from the microgrooves to the axonal terminals, or the longest length observed in the image. The tumor spheroid surface was determined using Imaris (Imaris x64 v8.3.1).

### Media compartmentalization and diffusion

Adjusting the media volumes in the soma and axonal compartments allows for the modification of hydrostatic pressure, enabling fluidic isolation in one compartment (compartmentalization) (Park et al., 2006; Taylor et al., 2003). The increased hydrostatic pressure resulting from the channel height overcomes the microfluidic resistance in the microgrooves. Consequently, matching the media volumes between compartments results in media diffusion through the microgrooves.

Compartmentalization or diffusion of media were tested by adding 3 KDa of 680-Alexa Fluor Dextran (Invitrogen, Cat#D34681) in the axonal compartment of DACIT, followed by collecting 3D stacks every hour over two days. Time-lapse images were analyzed using Fiji (Imagej2 version 2.14.0/1.54f). Briefly, the intensity decay of maximum intensity projection was corrected for photobleaching. Identical regions of interest were selected in the neuronal and the axonal compartments and the mean intensity signals were plotted over time.

## Results

### DACIT layer design and the fabrication of SU-8 master

We designed DACIT as two macrochannels (1.5 mm in width), each ending with a 2 mm well for loading media and cells (**Figure 1A**). Macrochannels are connected by the microgrooves (5 µm in width), which are aimed to prevent cells, while allowing axons to cross the channels. Microgrooves are designed to have a height of 3 µm to allow only axons, but not neurons (or Schwann cells) from the primary cultures, or cancer cells, to move across. At the same time, the height of the macrochannels needs to be sufficient to embed cancer spheroids (>200 µm) in ECM and allow for their growth and invasion in all three dimensions. To achieve such different heights, we have separated the device design into two layers and fabricated one mask containing the microgrooves (blue, **Figure 1B**) and another containing the macrochannels (red, **Figure 1C**). During the master fabrication, the designed layers, are overlaid (**Figure 1D**) using alignment marks (blue and red squares) present in both layers.

**Figure 1.**
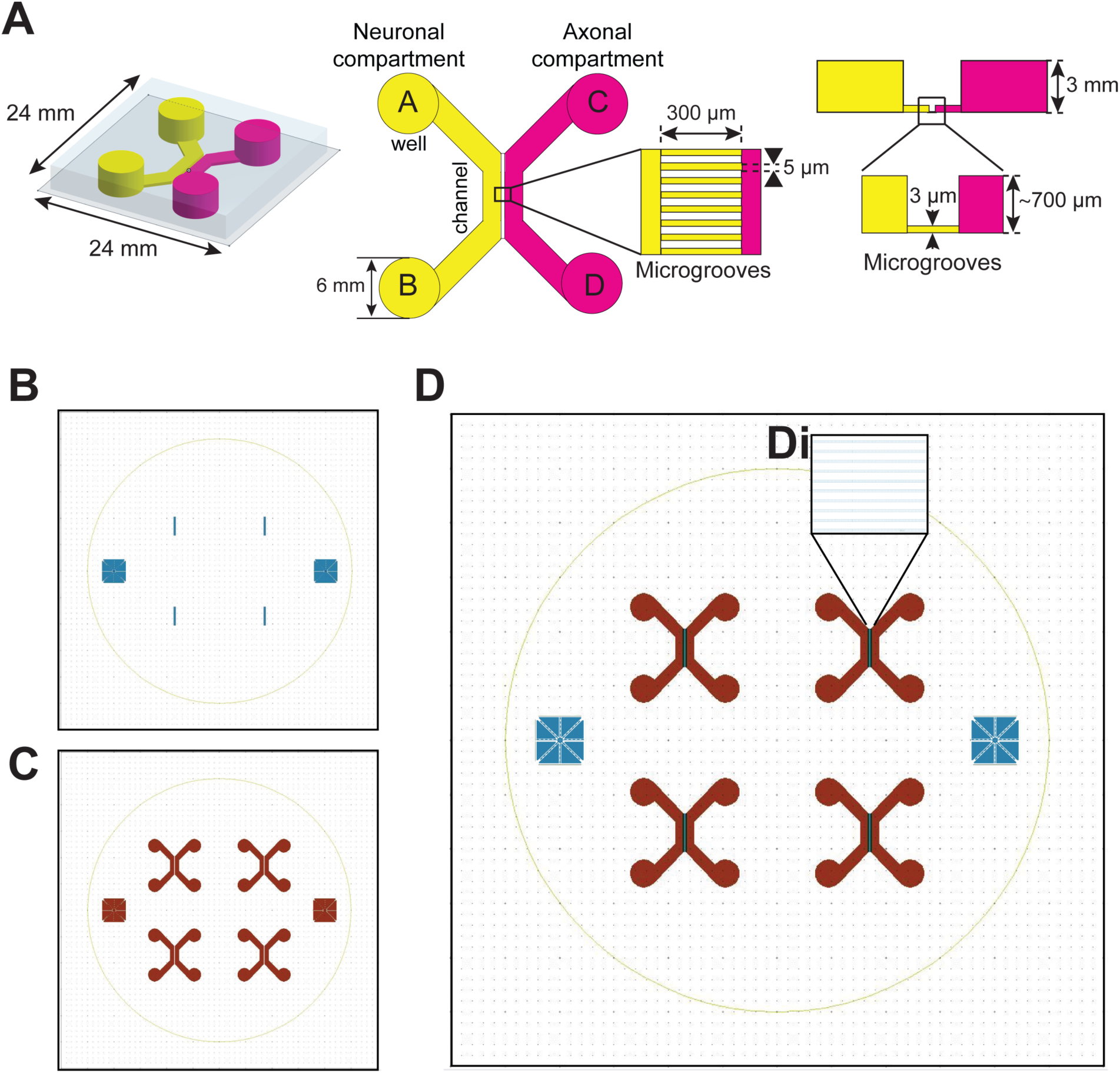
Design and alignment of the DACIT layers. **(A)** DACIT design in 3D view (left), top view (middle), and side view (right panel). Neuronal (yellow) and axonal (magenta) compartments contain two wells each, where solutions can be added (*A, C*) or removed (*B, D*). (**B**) Microgrooves layer with blue alignment marks on each side (**C**) Macrochannels and wells layer, with red alignment marks. (**D**) The overlay of layers for microgrooves (blue) and macrochannels (red); (**Di**) a closeup of the microgrooves connecting the left and the right macrochannel.

We used a multilayer strategy with the SU-8 photoresist 3000 resin to obtain the DACIT master (**Figure 2A**). Master contains four equal patterns and can produce four DACITs simultaneously (**Figure 2B**), using standard PDMS microfabrication protocols, punching out the wells and bonding PDMS to glass coverslip (Perrin et al., 2021). After producing final DACITs in PDMS, we have imaged the cross-section in z to measure the microchannel and confirmed a height of 700 µm (**Figure 2C**). As DACITs are mounted on 24x24 mm coverslips (**Figure 1A**), most standard microscopes will require additional holder to place them on the objective for imaging. Towards this goal, we designed and 3D-printed plastic holders useful for imaging one or six DACITs at a time (**Figure 2D**). For time-lapse imaging of DACITs, we have also CNC machined the six-DACIT holder out of aluminum (**Figure 2E**), which has a high thermal conductivity and allows maintaining the DACIT media temperature at 37 C° within the environmental chambers.

**Figure 2.**
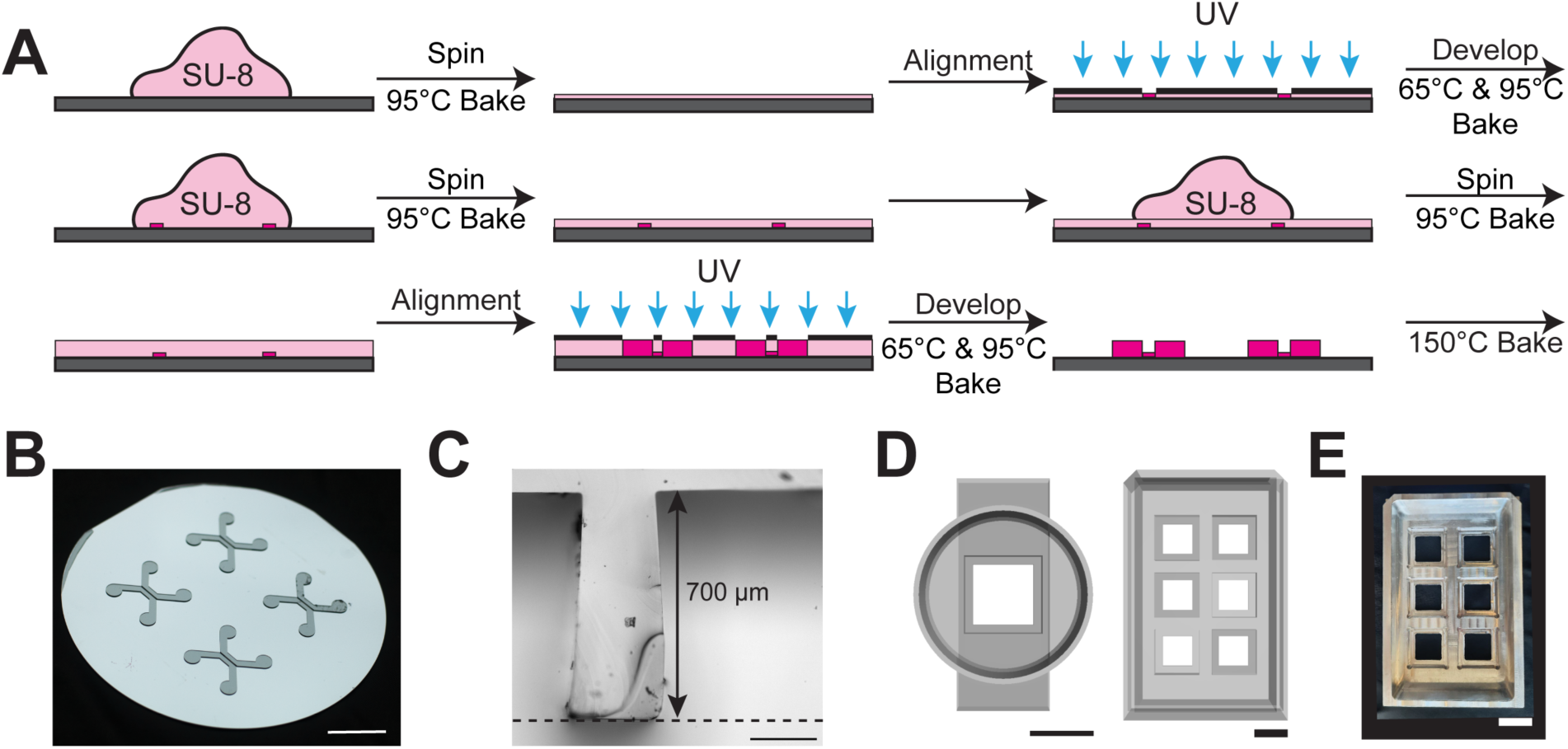
DACIT fabrication. (**A**) Main steps in the preparation of the SU-8 master include three rounds of spinning, mask alignment, UV exposure, and a combination of soft and hard-baking at 65, 95 and 150°C. (**B**) The final look of the SU-8 master. Scale bar: 2 cm. **(C**) A brightfield image of the DACIT cross-section shows that the channel height measures 700 microns. The dotted black line represents the position of the coverslip. Scale bar: 200 µm. (**D**) 3D-printed holders designed to mount and image either a single DACIT device (right) or six devices at a time (left). Scale bar: 2 cm. (**E**) Aluminum holder for 6 DACIT time-lapse imaging. Scale bar: 2 cm.

### DACIT assures the separation of cells and media between the two compartments

Some uses of DACIT will benefit from media mixing i.e. free diffusion, while for other uses, such as inhibitor treatments aimed at individual cell type, media should ideally remain within the assigned compartments. For example, to facilitate and speed up neoaxonogenesis, neuroscientists often use gradients of NGF (Lowry Curley et al., 2014), which would be ideal use for diffusion approach. However, culturing media for neurons and cancer cells are different, and in fact, serum used for cancer cell cultures can cause neurotoxicity (Brewer, 1998). Additionally, to test treatments with inhibitors or cancer cell drugs in a physiologically relevant fashion, we would like to have media limited to the axonal compartment. If media is maintained at the same level in both macrochannels, it will diffuse across, but a slight difference in volumes will stop diffusion and keep media within the appropriate compartment (Taylor & Jeon, 2010). To test the diffusion of medium across the microgrooves, we mixed the medium with 3 kDa dextran, loaded it to the axonal compartment of DACIT, and collected images every hour over 48 hours, at 37°C. We limited the time of the measurement to 48 h, since this is approximately the maximum time that cultures can be left without adding or changing the media. Following the compartmentalization approach, where the compartments are loaded with different volume of media (200 µl versus 150 µl, respectively), media permanently remains within its original compartment (**Figure 3A**, left). In contrast, using the diffusion approach, dextran diffuses across the microgrooves reaching an equilibrium over the first 6 hr (**Figure 3A**, right panel).

**Figure 3.**
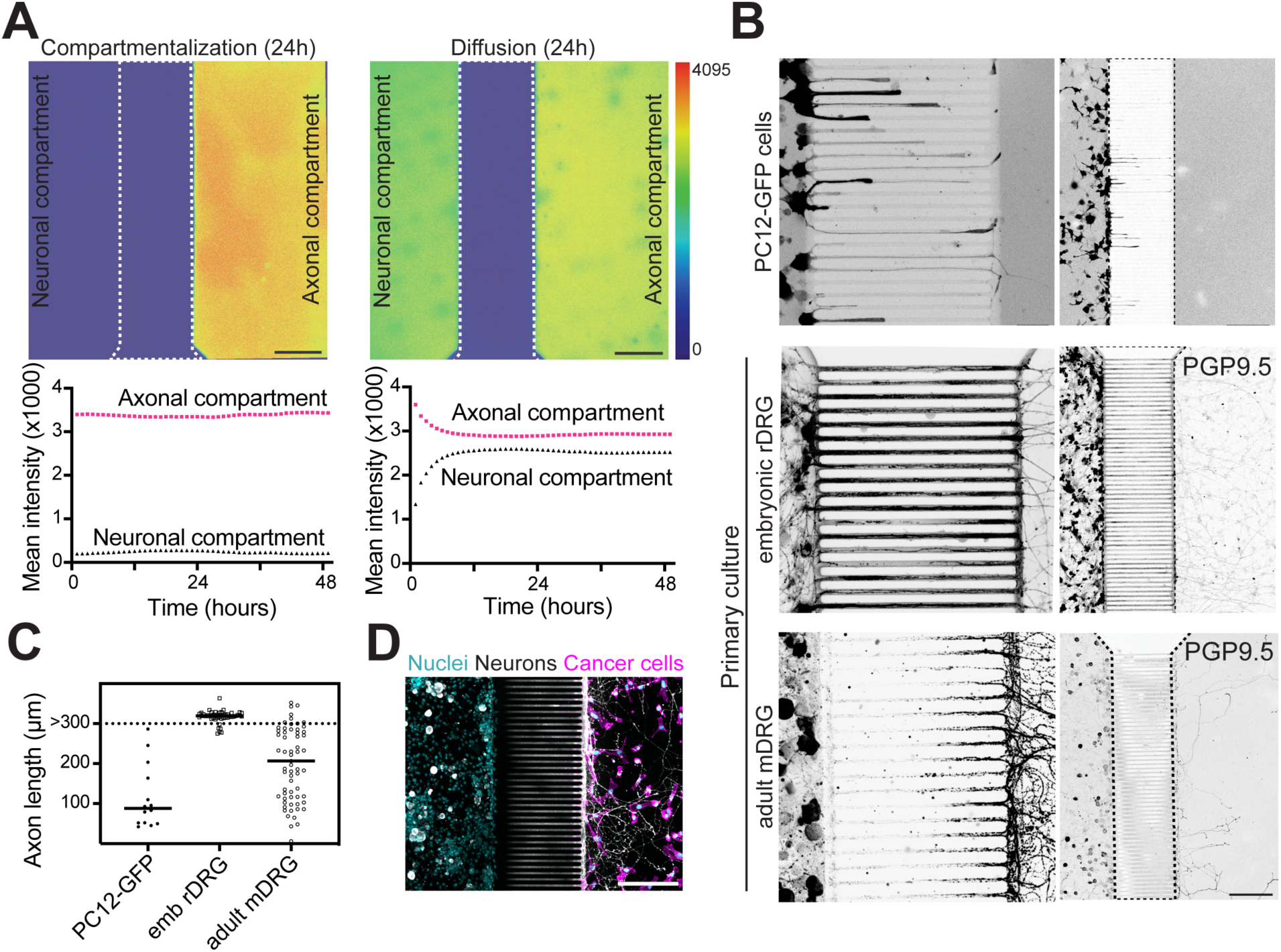
DACIT supports cellular separation and allows for choice of media compartmentalization or diffusion. Medium compartmentalization (**A**, left panels) or diffusion (**A**, right panels) was quantified using 48 h time-lapse recording of 3 kDa fluorescent dextran diffusion from the axonal compartment across the microgrooves. Pseudo-color images (**A**, top panels, look up table on the right) are stills showing the distribution of the dye at 24 h. Bottom panels show distribution of 3 kDa dextran over time (**A**, bottom panels). Axonal compartment readings are shown as red dotted lines and neuronal compartments as black dotted line. The scale bars equal 200 µm. (**B**) Images of PC12-GFP or PGP9.5 labeled primary DRG cultures, fluorescence is pseudo-colored black for clarity. A close-up view of axons extending across the microgrooves (**B**, left panel, Scale bar: 50 µm) and a larger view showing how axons populate the axonal compartment (**B**, right panel. Scale bar: 200µm). (**C**) Quantification of the axonal length from PC12 and primary DRG cultures. (**D**) Immunofluorescence of neuronal somas and axons (white, anti-PGP9.5) and cancer cells (magenta, F-actin and cyan, DAPI) interacting with axons in the axonal compartment. Scale bar:100 µm.

During DACIT design, we strived to optimize the length of the DACIT microgroove’s (300 µm) to ensure that most axons reach the axonal compartment while maintaining cancer cells and neuronal soma within their respective compartments. To test the timeline and efficiency of axon elongation across the microgrooves in DACIT in several widely-used models for neurons, we have compared the primary culture of adult murine DRGs, embryonic murine DRGs, and differentiated murine PC12 cells. Axon number, bundling and length vary depending on the number and type of neurons. However, in all cases, somas stay in the neuronal compartment and only axons reach the axonal compartment (**Figure 3B**). While PC12 cells are immortal and easy to culture and passage and represent sympathetic-like neurons (Chelmicka-Schorr et al, 1989), they grew relatively short neurites that did not always reach the axonal compartment (**Figure 3B**, top panel and **Figure 3C**). In contrast, most of the axons from the primary DRG culture successfully extended into the axonal compartment (**Figure 3B**, medium, and bottom panels and **Figure 3C**). As expected, embryonic axons extended faster, bundled more, filling the entire microgroove space, ended longer and more branched compared to those from adult animals (Rizk et al., 2023). Importantly, cellular separation was maintained over the course of the experiment, even after cancer cells were added to the axonal compartment and imaged at 96 hours (**Figure 3D**). Compared to previously published device, which demonstrated cancer cell migration across the microgrooves (Lei et al., 2016), DACIT is better suited for testing long-term cancer cell interactions with axons.

### DACIT measurements in neuronal and axonal compartment

DACIT offers a platform to evaluate neuronal or cancer cell behaviors following neoaxonogenesis in the axonal compartment (**Figure 4A**), while preserving anatomically relevant compartmentalization of cell bodies and media. Pharmacological treatments can be applied on a single, or both compartments simultaneously. DACIT enables the assessment of cancer cells’ ability to attach and spread, proliferate, migrate, and invade ECM in 2D or 3D environment, while maintaining their interaction with axons.

**Figure 4.**
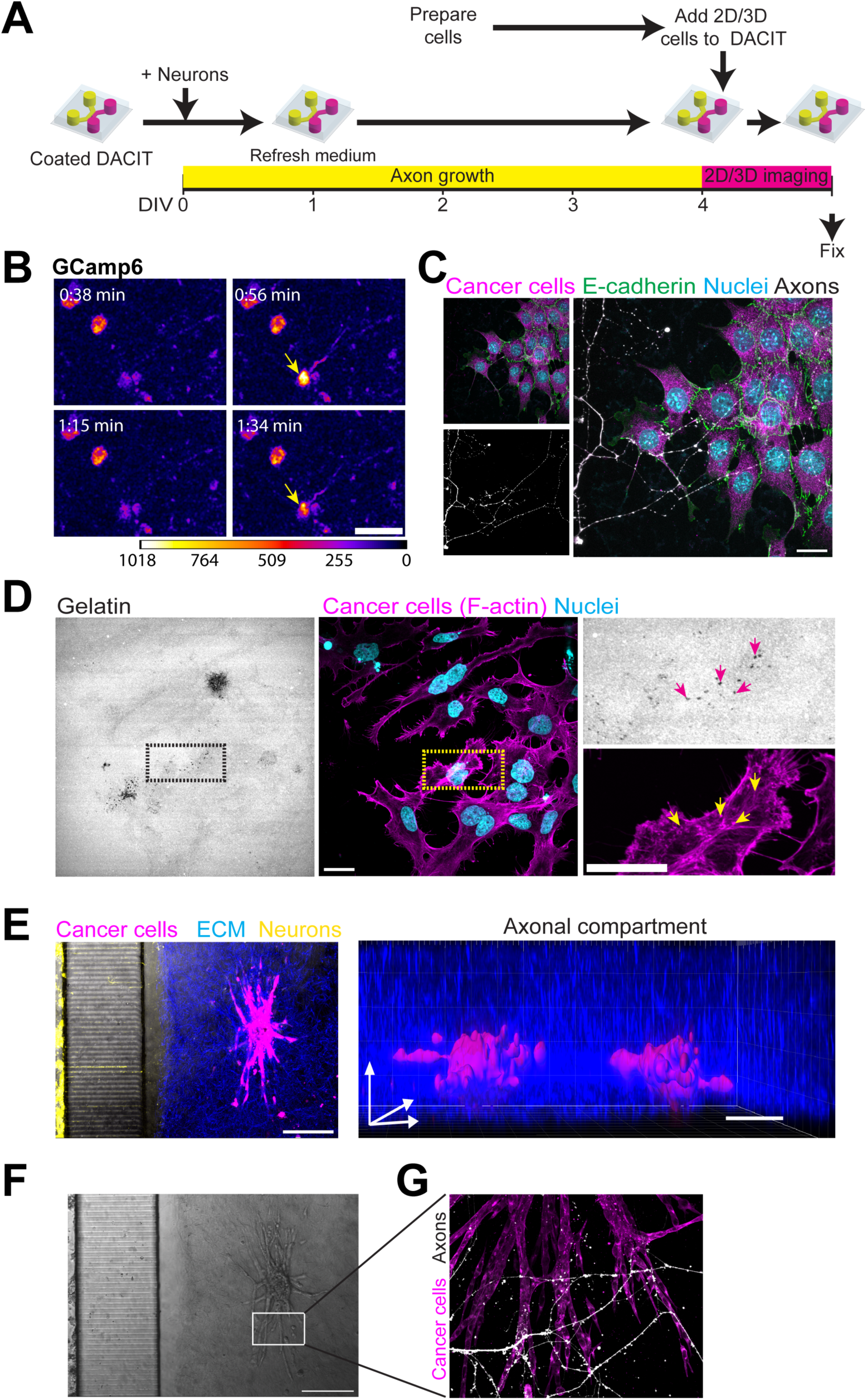
Invadopodia (2D) and spheroid (3D) invasion in DACIT. (**A**) Experimental timeline of 2D/3D invasion assays in DACIT. (**B**) Stills from a time-lapse recording of GCaMP6f-expressing sensory neurons cultured in the DACIT neuronal compartment (pseudo-color look up table on the bottom). Yellow arrows point to the cell exhibiting calcium transients (0.56 and 1:34 min). Scale bar = 50 µm. (**C**) Cancer cells (magenta, F-actin; DAPI, nuclei, blue; E-cadherin, green) plated on 2D gelatin/laminin ECM interact with axons (white, anti-PGP9.5). Scale bar: 25 µm. (**D**) Gelatin degradation puncta (black) generated by invadopodia (magenta, F-actin). Dashed boxes outline the zoomed-in area in right panels: invadopodia (yellow arrows, top) colocalize with gelatin degradation (magenta arrows, bottom). Scale bar: 25 µm. (**E**) 4T1-mScarlet spheroid (magenta) embedded in 3D ECM 405-collagen I: Matrigel mix (blue) interacting with neurons (yellow, PGP9.5) and overlaid with brightfield to demonstrate microgrooves (left panel); 3D Imaris reconstruction of two neighboring spheroids (right). Scale bars 200 µm. (**F**) Brightfield image of spheroid from (E). Note microgrooves on the left. Scale bar 200 µm. (**G**) Zoom-in to invasive strands of the spheroid (magenta, F-actin) interact with PGP9.5^+^ axons (white). Scale bar: 100 µm.

To demonstrate calcium dynamics in the neuronal compartment, we recorded time-lapses of GCaMP6f-sensory neurons (**Figure 4B** and **Supplementary Movie 1**). In the axonal compartment, time-lapse imaging of cancer cell attachment and migration in the presence of axons was monitored over 24h (**Supplementary Movie 2**). In **Figure 4C**, axonal compartment was fixed at 24 hours, showing that all cancer cells (magenta, F-actin and nuclei, cyan) are in physical contact with axons (PGP9.5, white).

To image invasion of cancer cells in the presence of axons in 2D, we modified the classical invadopodia assay, coating the axonal compartment with fluorescent gelatin first, followed by PLL and laminin. Once neurons were loaded and neoaxonogenesis occurred in the axonal compartment, cancer cells were plated. After 24 hours, at day 5 (**Figure 4A**), we were able to observe F-actin puncta characteristic for invadopodia precursors (**Figure 4D**, yellow arrows) colocalizing with degraded gelatin (**Figure 4D**, magenta arrows). In the 3D spheroid invasion assay, cancer cell spheroids were embedded in the collagen I: Matrigel mix and loaded in the middle of the axonal compartment. After 24 h, spheroids invaded into the matrix (**Figure 4E-G**), with invading strands in direct contact with axons (**Figure 4G**).

## Discussion

Recent studies have shown that the peripheral nervous system interacts with tumor and affects tumor progression (Ayala et al., 2001; Prazeres et al., 2020; Pundavela et al., 2015). However, studying the interaction between cancer cells and axons in a controlled system has proven to be challenging. Due to the absence of appropriate peripheral neuron models, most *in vitro* studies currently use primary cultures. Previous approaches have involved co-culturing cancer cells with dissociated (Le et al., 2022) or intact ganglia (He et al., 2015; Ni et al., 2020). However, non-neuronal cells present in these primary cultures, such as Schwann cells, may impact cancer cell behavior (Deborde et al., 2016; Deborde & Wong, 2022), and are difficult to remove from co-cultures while maintaining the viability of neuronal culture.

Here, we present a microfluidic device for axon-cancer cell interaction (DACIT) in 2D and 3D. DACIT grants a separation between the two cellular compartments: neuronal compartment, able to extend axons across the microgrooves, and the axonal compartment, in which cancer cells are loaded and interact with axonal terminals. Importantly, DACIT’s dimensions minimize reagent waste and maximize the use of precious primary culture samples with small cell numbers. Here, we have demonstrated the utility of DACIT by using either dissociated embryonic or adult murine DRGs in the neuronal compartment, and murine cancer cell lines plated in 2D or as 3D spheroids embedded in ECM in the axonal i.e. cancer cell compartment. However, the versatility of this device is not limited to the murine cells. Neuronal compartment of DACIT can also be loaded with intact DRGs (murine or patient-derived) or human neurons derived from the induced pluripotent stem cells (iPSCs), while the axonal (cancer cell) compartment can host non-cancer cells or patient-derived cancer organoids, setting the stage for drug efficiency testing in translational or clinical research.

When using unsealed microfluidic devices such as DACIT, performing timelapse imaging can be challenging due to the heat generated during imaging, which can cause faster evaporation. Therefore, it’s crucial to use imaging chambers that can maintain optimal humidity. Our custom 3D-printed holders are designed to be used with lids from the microplate or 35 mm culture dish, which helps to maintain humidity. For time-lapse imaging, we have also created a custom aluminum holder which allows maintaining temperature (**see Figure 2E**). To maintain humidity and reduce evaporation during long-term imaging, additional preventative measures can be taken: a layer of mineral oil can be added on top of the media, and open 35 mm dishes with PBS can be placed around the DACIT.

While axon bundles and even single axons are sometimes visible using brightfield (**Supplementary Movie 2**), axon labeling facilitates visualization and assures all axons will be appropriately segmented and quantified. To image live axons in DACIT, retrograde tracer dyes can be used (Cheng et al., 2014; Kallogjerovic et al., 2024b). Additionally, adeno-associated viruses (AAV) expressing fluorescent proteins under neuron-specific or ubiquitous promoters can be added to the primary culture several days prior to plating in DACIT (see **Figure 4B** and **Supplementary Movie 1**) (Gordon et al., 2013). Importantly, as protein expression requires multiple days and primary cultures cannot survive long-term, experiments inducing the expression of fluorescent proteins via AAVs should be carefully planned.

Using compartmentalization or diffusion loading strategies, small molecules in DACIT either diffuse or remain in a specific compartment. The compartmentalization in DACIT prevents flow from the lower- to the higher-volume compartment, promoting the localized enrichment of neuron-secreted factors within a single compartment. Taking advance of this, we showed that DACIT can be used to retrograde trace sensory neurons (Kallogjerovic et al., 2024b)

To bond PDMS devices to coverslips, Poly-L-Lysine (PLL) is often used, which allows detachment of PDMS following plating neurons, facilitating analysis of fixed cells (Park et al., 2006; Taylor et al., 2005). For DACIT fabrication, we used oxygen plasma treatment to bond the PDMS with the glass coverslip. This approach permanently separates the compartments and allows for monitoring axon-cancer cell interactions over multiple days.

Behavior of both neurons and cancer cells depends on the ECM that cells adhere to (Harris et al., 2017). Common substrates that favor neuron attachment and axonogenesis are PLL followed by laminin or Matrigel. For invadopodia assays, the conditions for visualization and measurement of invadopodia-based ECM degradation were perfected using fluorescent, cross- linked gelatin. To allow for proper axonogenesis in axonal i.e. cancer cell compartment without affecting invadopodia measurements, we overlaid the gelatin layer with PLL and laminin. In the case of 3D invasion assays, Matrigel was mixed with collagen I to promote axons to reach tumor spheroids. Notably, despite the high density of the 3D ECM or the presence of primary culture debris in the soma compartment, channel obstruction, frequently observed in microfluidic devices, is not an issue in DACIT, likely due to the sufficiently high of the macrochannels.

One of our main goals in developing DACIT was to perform 3D cell culture assays that better recapitulate cancer cell interactions, proliferation, and resistance *in vivo* (Åkerlund et al., 2023; Lv et al., 2017). Fabricating a SU-8 master mold with a high aspect ratio is a challenging task in lithography due to the possibility of collapse (Amato et al., 2012). Several approaches, such as e-beam, X-ray, and two-photon lithography, as well as optimizing post-exposure and soft-baking temperatures, have been developed to address this problem. However, these approaches are often associated with drawbacks such as high costs, time consumption, and low temporal stability (Anhoj et al., 2006; Cadarso et al., 2010; Del Campo & Greiner, 2007; Keller et al., 2008; Lin et al., 2002).

To create a device which simultaneously isolates axons from their cell body (3 µm-high microgrooves) and allows for multi-day growth and invasion of 3D tumor spheroids or organoids (700 µm-high side compartments), we used two consecutive spinning steps of UV-lithography with SU-8 3000 epoxy resin (Zhou & Huang, 2018). Since 3 µm resin is thinner than the standard working range of SU-8 photoresists (Kayaku Advanced Materials,. Inc), while, at the same time, 700 µm is thicker than any other reported device, we systematically tested a range of combinations for the baking temperature and duration, as well as the spinning ramp and speed. Creating a 700 µm-thick macrochannel thickness required applying SU-8 twice and fine-tuning process parameters. The most critical step in making 700 µm-thick chambers was to align the photomask for exposure with the already patterned microgrooves’ layer. The technical challenge of this step lays in the possibility of shearing the uncured layers due to the attachment of the photoresist to the photomask. Successfully produced master was multiplied using Smooth-On mold, to allow for production of up to 20 DACIT devices simultaneously.

In this work, we demonstrated that DACIT facilitates visualization and manipulation of the interactions between axons and cancer cells in a physiologically relevant context. We show that DACIT can be successfully used to study the interactions between wide range of models of peripheral neurons and cancer cells in 2D and 3D. Measurements of the neuronal effect on the cancer cell adhesion, morphology, migration, and invasion, as well as cancer cell effect on the neoaxonogenesis of different neuronal subtypes and neuron activity. In the future, DACIT can be easily adapted to test drug effects on interactions between neurons and cancer cells, or even to analyze the effect of peripheral neuron interactions with cell types other than cancer cells, including immune, endothelial, or muscle cells. Moreover, human induced pluripotent stem cell (hiPSCs) differentiation into functional peripheral neurons has been described. (Fujimori et al., 2018; Hiranuma et al., 2024a; Schwartzentruber et al., 2017). In the future, this class of neuron models can be combined with 3D patient-derived organoids in DACIT for studying neuron-cancer cell interactions in human cells.

## Acknowledgments

We thank the following funding sources for their support: NIH NCI R01 CA230777; American Cancer Society Research Scholar Grant 134415-RSG-20-34-01-CSM, DOD BCRP Breakthrough Award BC230197, PA-CURE and WW Smith Charitable Trust awards to B.G. and METAvivor Early Career Research Award to I.V.Q. This work was carried out in part at the Singh Center for Nanotechnology, part of the National Nanotechnology Coordinated Infrastructure Program, which is supported by the National Science Foundation grant NNCI-2025608. We also thank Dr. Eric Johnston from Nanotechnology Singh Center for his help with fabrication of the SU8 master, Fox Chase Cancer Center Machine shop, and Peter Lelkes lab for the kind gift of PC12-GFP cells.

## Author contributions

Conceptualization: I.V.Q., V.A., E.T., B.G.; Sample preparation: I.V.Q., K.A., V.A., E.B., N.H., X.Z, S.K. Data acquisition, analysis, and interpretation: I.V.Q., K.A., V.A., E.B., S.K., B.G.; Writing: I.V.Q., K.A., V.A., E.B., E.T., N.H., X.X., G.T. and B.G.; Supervision: E.T., G.T. and B.G.

## Supplementary files

**Supplementary Movie 1. GCaMP6f dynamics in DACIT plated-sensory neurons.** Time-lapse of GCamMP6f-neurons exhibiting spontaneous calcium transients in the neuronal compartment of DACIT (recorded at 14 Hz for 1:45 min). Scale bar 50 µm.

**Supplementary Movie 2. Axons interacting with 4T1 cells on DACIT.** Time-lapse recording of 4T1 cells spreading in the presence of axons, in the axonal compartment of DACIT (recorded every 10 minutes for 20 hours). Scale bar 50 µm.

